# Osteosarcoma with apparent Ewing sarcoma gene rearrangement

**DOI:** 10.1101/039834

**Authors:** Melissa Mathias, Alexander J Chou, Paul Meyers, Neerav Shukla, Meera Hameed, Narasimhan Agaram, Lulu Wang, Michael F. Berger, Michael Walsh, Alex Kentsis

## Abstract

Poorly differentiated round cell sarcomas present diagnostic challenges due to their variable morphology and lack of specific immunophenotypic markers. We present a case of a 15-year-old female with a tibial tumor that exhibited features of Ewing-like sarcoma, including apparent rearrangement of the *EWSR1* gene. Hybridization capture-based next-generation DNA sequencing showed evidence of complex genomic rearrangements, absence of known pathogenic Ewing-like chromosome translocations, and deletions *RB1*, PTCH1, and *ATRX*, supporting the diagnosis of osteosarcoma. This illustrates the potential of clinical genomic profiling to improve diagnosis and enable specifically targeted therapies for cancers with complex pathologies.

## INTRODUCTION

Recent introduction of molecular and genomic profiling in clinical oncology has transformed our understanding and treatment of many human cancers. In particular, though classic descriptions of bony tumors have included osteogenic and Ewing sarcomas, recent studies have begun to reveal subtypes of round cell sarcomas with shared pathogenic mechanisms arising in diverse tissues, as well as distinct tumor subtypes with similar appearance. For example, classic Ewing bone sarcomas, extraosseous Ewing-like sarcomas, desmoplastic small round cell tumors of the abdomen, and extraskeletal myxoid chondrosarcomas have diverse clinical presentations and histologic morphologies, but all share characteristic translocations involving the Ewing sarcoma breakpoint region 1 (*EWSR1)* gene and molecular pathogenic mechanisms with poor responses to conventional chemotherapy ^1,2^. Conversely, round cell variants of osteosarcoma, Ewing sarcoma, peripheral neuroectodermal tumor, and myeloid leukemia can have indistinguishable morphologic appearance and share immunophenotypic markers, in spite of distinct molecular and genetic pathologies ^3^. In particular, osteosarcomas are characteristically defined by complex genomic rearrangements, including chromothripsis, and inactivating mutations of *TP53* and *RB1* genes ^4^. On the other hand, most Ewing sarcomas are genomically stable with low rates of aneuploidy and chromosomal copy number changes, infrequent mutations of *TP53*, and uniform chromosomal translocations of the *EWSR1* gene ^5-7^. Cases of small cell osteosarcomas with *EWSR1* rearrangements have been reported, but it is not known whether this is associated with Ewing-like sarcoma clinical behavior and therapy response, a question with potential therapeutic implications since treatment regimens for osteogenic and Ewing sarcomas have significant differences ^8^. Here, we describe a patient with a tibial tumor, who was found to have a round cell sarcoma with an apparent *EWSR1* rearrangement, consistent with a diagnosis of Ewing-like sarcoma, but instead was diagnosed as osteosarcoma based on genomic profiling.

## CASE DESCRIPTION

The patient is a 15-year-old female with an unremarkable past medical history who developed a slowly enlarging painful tibial mass. Radiography and magnetic resonance imaging revealed a heterogeneous mass in the metadiaphysis with periosteal reaction and extensive soft tissue extension. Plain radiography described the lesion as containing mixed lytic and sclerotic lesions. Computed tomography described peripheral sclerosis extending into the medial tibial epiphysis. Her family history was significant for two maternal great aunts with breast cancer in their fifties and sixties, a maternal great grandfather with lung cancer, and a paternal grandfather with prostate cancer.

Percutaneous tumor biopsy revealed large round cells with clear to granular cytoplasm, and focal areas of collagen matrix. Immunohistochemical analysis of this biopsy specimen demonstrated tumor cells that were diffusely positive for membranous CD99 and focally positive for CD68, but negative for cytokeratin, S100, HMB45, A103, LCA, desmin, SATB2, and EMA. Fluorescence in situ hybridization (FISH) analysis using Vysis probes (Abbott) demonstrated *EWSR1* gene rearrangement in 56% of the cells in a focal area of the tumor (Figure 1). The remaining tumor in the biopsy showed 3-4 copies of *EWSR1* gene in 74% of the cells. Given the focal nature of *EWSR1* rearrangement and recent discovery of round cell Ewing-like sarcomas with *CIC* gene translocations without *EWSR1* rearrangements ^9^, FISH for *CIC* rearrangements was performed and found to be normal (not shown). Similarly, FISH analysis using custom BAC probes showed no rearrangements of the *FLI1*, *ERG* or *NFATC2* genes ^10^. Based on these histopathologic findings, a diagnosis of Ewing-like sarcoma was made, and neoadjuvant Ewing sarcoma-directed chemotherapy treatment was initiated with high-dose cyclophosphamide, doxorubicin, and vincristine ^11^.

**Figure 1:**
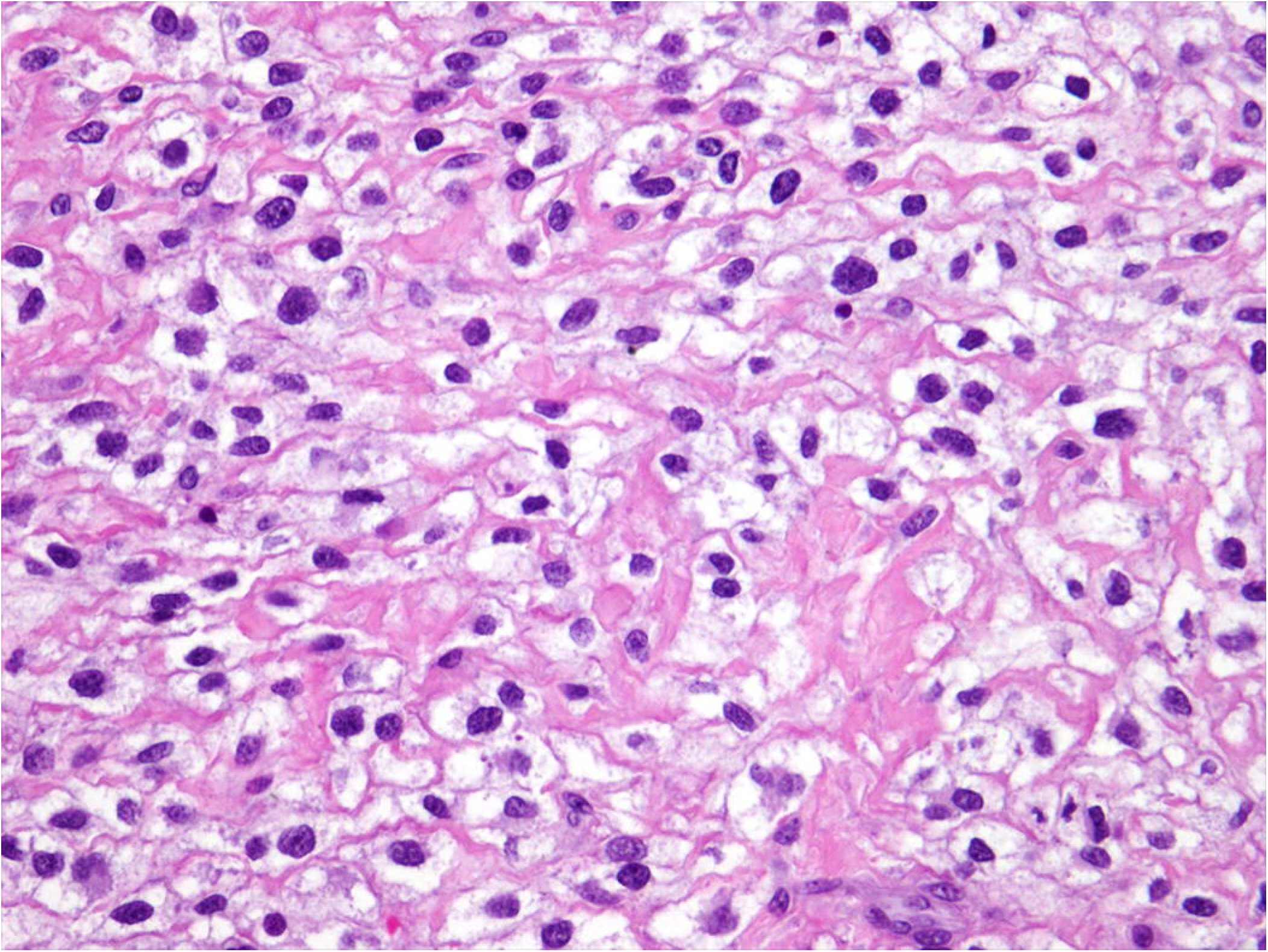
Round cell sarcoma with apparent EWSR1 rearrangement: Percutaneous biopsy section stained with hematoxylin and eosin, demonstrating monomorphic round cell sarcoma cells

**Figure 2:**
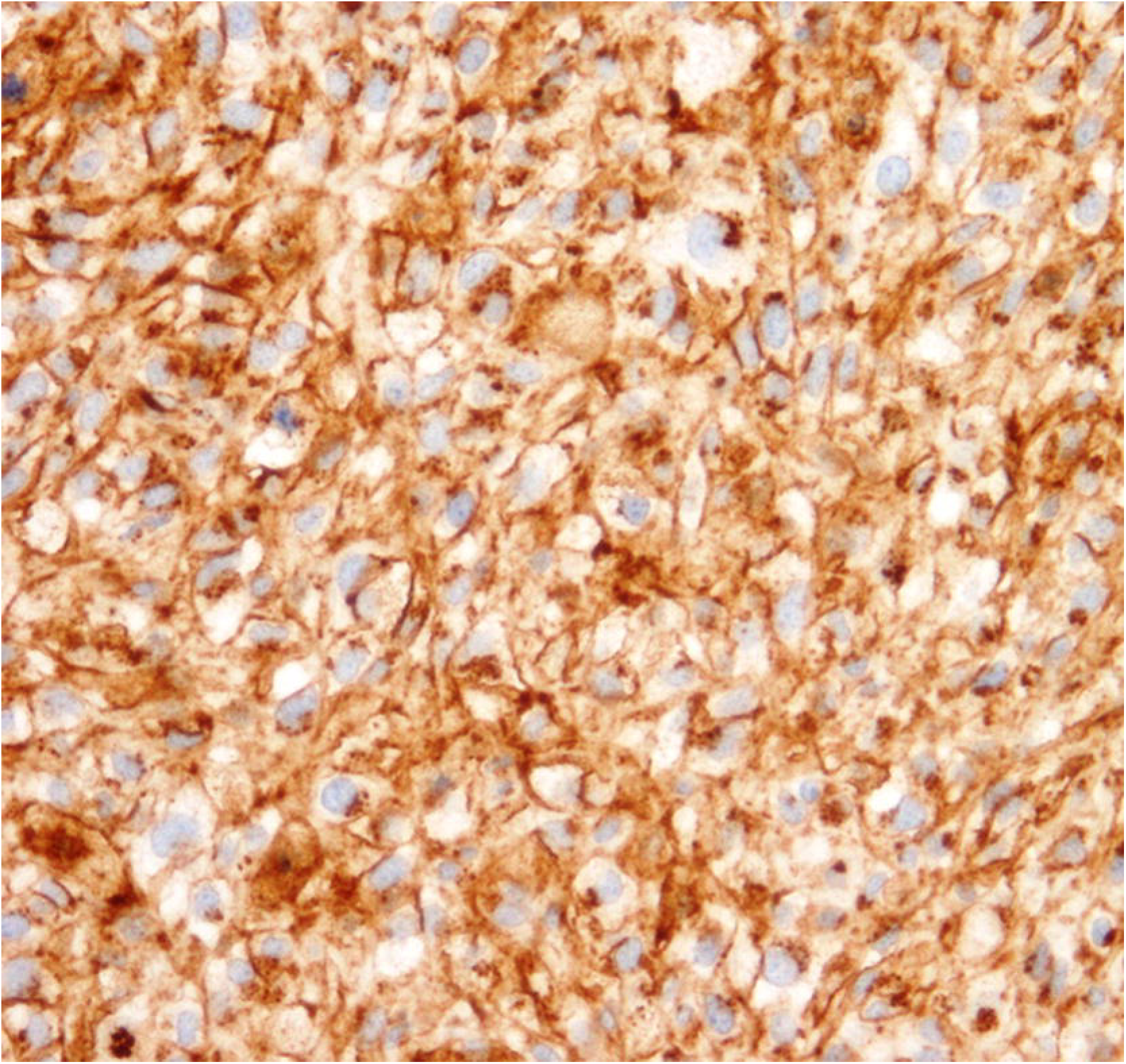
Immunohistochemistry, demonstrating diffuse membranous CD99 staining

**Figure 3:**
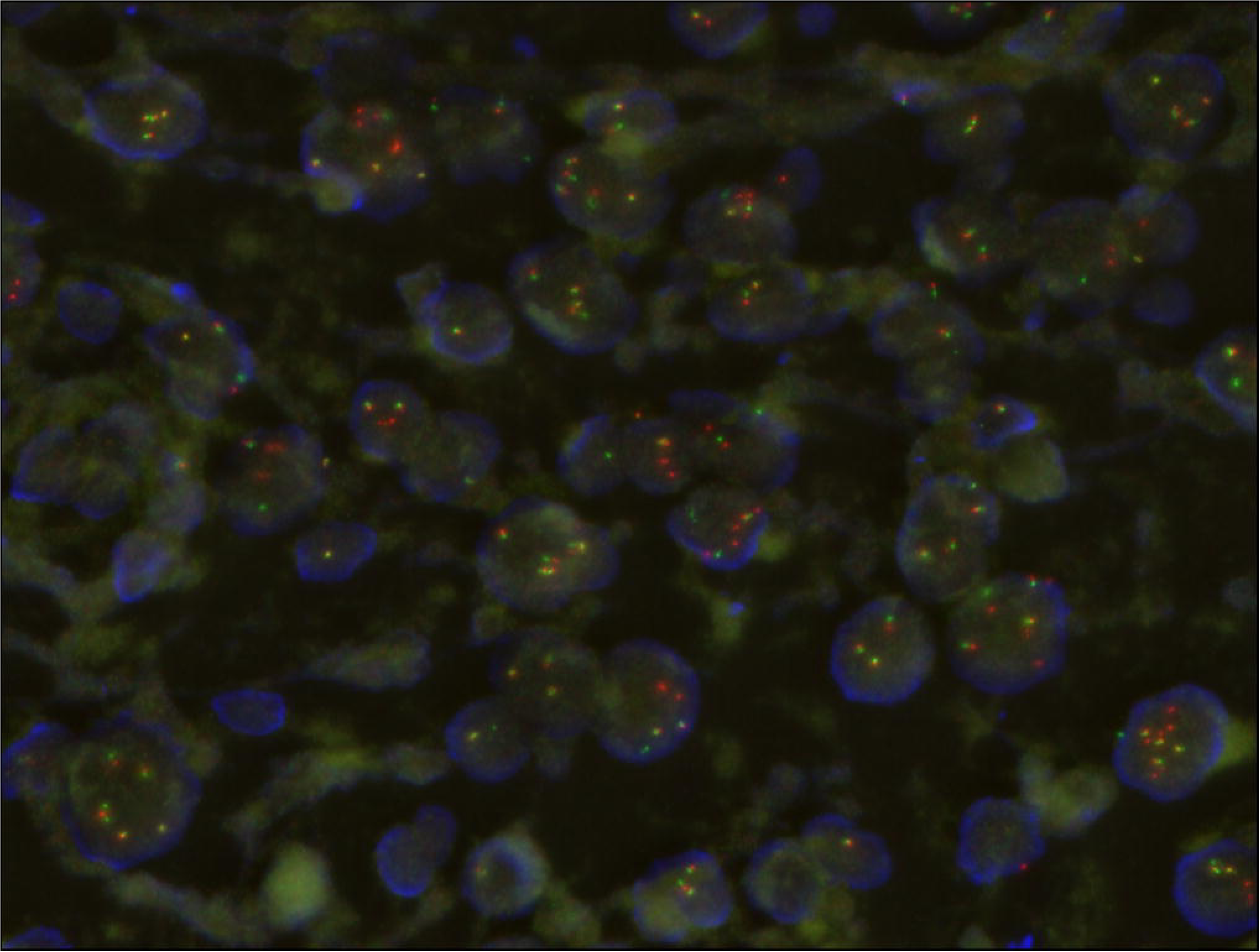
Fluorescence in situ hybridization (FISH) micrograph, demonstrating rearrangements of the 5’ and 3’ regions of the *EWSR1* gene labeled in red and green, respectively, with occasional yellow signals denoting intact *EWSR1* genes. Nuclear DNA is labeled in blue.

Given the atypical histopathologic tumor appearance and atypical FISH findings, the tumor specimen and matched peripheral blood cell genomic DNA were analyzed using hybridization capture-based next-generation DNA sequencing, specifically designed to identify mutations in 341 genes known to be pathogenically mutated in human cancers ^12^. These results are detailed in Table 1. Notably, we found numerous copy number changes, including homozygous segmental chromosomal deletions involving *RB1, PTCH1*, and *ATRX*, genes that are characteristically disrupted in osteosarcomas instead of Ewing sarcomas ^13-15^. Additionally, the analysis demonstrated absence of genomic rearrangements affecting specific regions of the *EWSR1* gene known to be involved in Ewing-like chromosomal translocations ^12,16^. The high number of chromosomal alterations seen in this patients tumor is consistent with chromoplexy classically seen in osteosarcoma ^13,15^.

Given the presence of i) numerous genomic copy number changes, ii) mutations of genes characteristically disrupted in osteosarcomas, and iii) lack of Ewing-like chromosomal rearrangements, the patient’s original diagnosis was reassessed. While genomically complex Ewing sarcomas have been observed, they are invariably associated with mutations of *TP53* ^5^. In this case, we found no pathogenic somatic or germ-line alterations of *TP53*. However, subsets of osteosarcoma have been found to exhibit *TP53* rearrangements involving *Alu* intronic elements, which may evade detection by targeted sequencing such as MSK-IMPACT ^12,17,18^. At the time of this reassessment, the patient underwent definitive surgical resection, which demonstrated high-grade osteosarcoma with 80% cellular necrosis corroborating the conclusions of the molecular analysis. Given the definitive diagnosis of osteosarcoma, the patient completed consolidation therapy with osteosarcoma-directed treatment using high-dose methotrexate, doxorubicin, and cisplatin ^11^. Consistent with the patient’s remote family history of cancer, examination of patient‘s non-tumor DNA revealed no pathogenic germ-line variants. Variants of unknown significance in her germline included only PARP1 p.W481C, FOXP1 p.A15V, RECQL4 p.D1121N, and AMER1 p.E385Q.

## DISCUSSION

Poorly differentiated round cell sarcomas present significant diagnostic challenges due to their variable morphology and lack of specific immunophenotypic markers ^3^. Genetic tests are increasingly being used to improve both the classification of disease in general and precision of diagnosis for individual patients. In this case, the diagnostic tumor specimen exhibited an apparent *EWSR1* gene rearrangement and CD99 immunopositivity consistent with Ewing-like sarcoma. However, in addition to Ewing sarcomas, CD99 expression can be seen in synovial sarcomas and small cell osteosarcomas ^8^. This case suggests that additional variants of osteosarcomas appear to express CD99. In addition, genomic instability and chromothripsis that characterize distinct tumors such as osteosarcoma can induce apparent gene rearrangements that are in and of themselves pathogenic ^4,19^. Hybridization capture-based next-generation DNA sequencing targeting the specific region of *EWSR1* characteristically translocated in Ewing sarcomas demonstrated absence of known pathogenic Ewing-like translocations, and presence of numerous copy number changes consistent with genomic instability (Table 1) ^12^. We suspect that similar genomic rearrangements are responsible for the apparent *EWSR1* break-apart FISH signal (Figure 1B). In contrast to focused clinical genetic tests such as FISH, genomic profiling using next-generation DNA sequencing offers the ability to characterize genetic lesions genome-wide, with base-pair resolution necessary to confidently ascribe pathogenic functions. Indeed, clinical genomic profiling utilized in this case revealed deletions of genes characteristically inactivated in osteosarcomas, such as *RB1*, *ATRX*, and *PTCH1* ^13,14^. Thus, genomic profiling enhanced clinical diagnosis, leading to more precise therapy selection, given substantial differences in current treatment regimens for osteogenic and Ewing sarcomas ^11^. In this specific case, we identified somatic deletion of *PTCH1*, consistent with the prior observations of *PTCH1* mutations in osteosarcomas, which cause activation of the Hedgehog signaling pathway and susceptibility to emerging targeted inhibitors of Hedgehog signaling such as vismodegib ^20^. We anticipate that comprehensive use of clinical genomic profiling will lead to the identification of pathogenic lesions with specific therapeutic agents allowing for their rapid incorporation into precisely engineered treatment regimens, particularly for cancers with complex or atypical pathologies, similar to the one described here.

## Acknowledgements

This work was supported by the NIH K08 CA160660 and the Burroughs Wellcome Fund (A.K).

## ADDITIONAL INFORMATION

### Data Deposition and Access

None.

### Ethics Statement

The patient and family gave assent and consent to all parts of this study.

### Author Contributions

Melissa Mathias had full access to all of the data in the study and takes responsibility for the integrity of the data and the accuracy of the data analysis. *Study concept and design*: Alex Kentsis and Melissa Mathias. *Acquisition, analysis, and interpretation of data*: All authors. *Drafting of the manuscript*: Mathias and Kentsis. *Critical revision of the manuscript for important intellectual content*: All authors. *Statistical analysis*: Not applicable. *Obtained funding*: Not applicable. *Administrative, technical, or material support*: Kentsis. *Study supervision*: Kentsis.

